# Alpha Event-related Desynchronization During Reward Processing in Schizophrenia

**DOI:** 10.1101/2021.02.25.432936

**Authors:** Tobias F. Marton, Brian J. Roach, Clay B. Holroyd, Judith M. Ford, John R. McQuaid, Daniel H. Mathalon, Susanna L. Fryer

**Affiliations:** Department of Psychiatry, Weill Institute for Neurosciences, University of California San Francisco, San Francisco, CA; San Francisco VA Healthcare System, San Francisco, CA; Department of Experimental Psychology, Ghent University, Belgium

## Abstract

**Background:** Deficits in the way the brain processes rewards may contribute to negative symptoms in schizophrenia. Synchronization of alpha band neural oscillations is a dominant EEG signal when people are awake, but at rest. In contrast, alpha desynchronization to salient events is thought to direct allocation of information processing resources away from the internal state, to process salient stimuli in the external environment. Here, we hypothesize that alpha event-related desynchronization (ERD) during reward processing is altered in schizophrenia, leading to less difference in alpha ERD magnitude between winning and losing outcomes.

**Methods:** EEG was recorded while participants (patients with schizophrenia (SZ)=54; healthy controls (HC) = 54) completed a casino-style slot machine gambling task. Total power, a measure of neural oscillation magnitude was measured in the alpha frequency range (8-14 Hz), time-locked to reward delivery, extracted via principal components analysis, and then compared between groups and equiprobable win and near miss loss reward outcomes. Associations between alpha power and negative symptoms and trait rumination were examined.

**Results:** A significant Group X Reward Outcome interaction (p=.018) was explained by differences within the HC group, driven by significant posterior-occipital alpha desynchronization to wins, relative to near miss losses (p<.001). In contrast, SZ did not modulate alpha power to wins vs. near miss losses (p>.1), nor did alpha power relate to negative symptoms (p>.1). However, across all participants, less alpha ERD to reward outcomes was related to more trait rumination, for both wins (p=.005) and near-miss losses (p=.002), with no group differences observed in the slopes of these relationships.

**Conclusion:** These findings suggest that event-related modulation of alpha power is altered in schizophrenia during reward outcome processing, even when reward attainment places minimal demands on higher-order cognitive processes during slot machine play. In addition, high trait rumination is associated with less event-related desynchronization to reward feedback, suggesting that rumination covaries with less external attentional allocation to reward processing, regardless of reward outcome valence and group membership.

## Introduction

Dysfunction within the neural systems involved in reward processing is a primary candidate mechanism underlying hedonic and motivational deficits in schizophrenia(1, 2). However, how and when impairments arise along the pathway between reward potential and reward attainment is still being characterized. Developing a more precise understanding of the neurobiology underlying reward system deficits in schizophrenia may help address the unmet need for interventions targeting negative symptoms(2). An extensive body of neurobehavioral data describes relatively intact reward sensitivity and hedonic experience (3), including implicit reward processing in patients with schizophrenia, suggesting areas of preserved reward functioning in schizophrenia (4). However, behavioral deficits emerge with downstream aspects of reward processing, such as representing reward values and reward-based learning (5–7). Data are converging across methodologies to suggest that altered reward processing in schizophrenia results in an inability to optimally use motivational information to guide behavior toward goal attainment (8). These findings suggest that in order to understand how reward-relevant network dysfunction ultimately translates to functional deficits in goal pursuit and attainment in schizophrenia, it will be important to elucidate the temporal dynamics of how reward deficits unfold, as well as how reward-related signals interact with the cognitive processes like attention and working memory to enable motivated approach behaviors (9). Electroencephalography-based measures of neural oscillations offer a compelling method to probe the temporal course of brain responses to reward (10).

Time-frequency based analyses can be used to parse EEG data into measures of neuro-oscillatory magnitude and phase. This information is of functional significance because distinct cognitive processes are thought to correspond to different “signatures” within the brain’s oscillatory structure (e.g., within canonical frequency bands delta, theta, alpha, beta, gamma), and synchronization within and across frequency bands is thought to constitute an elemental mechanism of information processing and binding within a functional neural network (11). The alpha band (~10 Hz) is a predominant rhythm in the human brain and has been studied in relation to numerous cognitive processes (12). A relative decrease in the magnitude of alpha oscillations in response to a task probe, referred to as event-related desynchronization (ERD), is elicited across a broad range of cognitive and sensorimotor tasks and is thought to reflect engagement of fundamental cognitive processes, such as attention and working memory, required for task engagement and appropriate response selection(13). Therefore, elucidating the temporal structure and topography of alpha ERD may provide insights into processes that support effective task engagement. Indeed, posterior alpha ERD in the context of visual processing is thought to relate to the functional activation of cortical networks relevant for extrinsic attention(14). In the context of a visual task, alpha ERD manifests as a stimulus-locked reduction in alpha power across the occipital lobe; this reduction in alpha power may reflect a “release” of inhibition in visual cortical areas allowing transition from an “off-line” to “on-line” information processing state in response to task engagement and salient stimulus presentation (12).

Attentional processes reflected by alpha ERD are modulated by many factors, including reward availability. For example, in an incentivized version of the paradigm in which Pfurtscheller and Aranibar originally defined alpha ERD, incentive (relative to non-incentive trials) elicited faster behavioral responses and widespread alpha ERD, with the fast reaction time trials showing the largest alpha ERD responses (15). Alpha ERD to target cues is enhanced (i.e., greater alpha suppression) by reward in a rapid visual attention task, an effect thought to be driven by the increased attentional drive in response to reward motivation (16). Similarly, an incentivized Stroop task revealed increased alpha ERD to cued reward relative to non-reward trials, that corresponded with faster reaction time.

Coordinated neural oscillatory activity is a fundamental organizing property of brain functioning, and alterations in these processes have been implicated in neuropsychiatric disease (17, 18). Neural oscillatory dysfunction has been described in schizophrenia and may underlie features of schizophrenia pathophysiology (for review, see (19, 20), including reductions in alpha band ERD(21–26). For example, reductions in both alpha and beta ERD in schizophrenia (SZ) (i.e., reduced suppression) has been demonstrated during working memory consolidation (26). Alpha ERD is also relevant to selective attention in the visual system, in which alpha oscillatory abnormalities have been observed in SZ, including elevated baseline (pre-stimulus) alpha as well as reduced visual stimulus-induced ERD. Within SZ, the degree of alpha ERD impairment correlated not only with target detection sensitivity (d’) during the visual task, but also with MATRICS Consensus Cognitive battery(27) attention/vigilance and visual learning domain scores, consistent with a top-down attentional mechanism modulating alpha desynchronization(25).

Indeed, recent conceptualizations posit that rather than being functionally distinct circuits, reward and attentional networks are reciprocally linked; Reward-centered salience and motivational pathways directly modulate prefrontal attentional circuitry, while “top down” cognitive processes mediated by the dorsal attentional network modulate reward sensitivity and reward-based learning (28). However, to date, few studies in SZ directly investigate the task engagement reflected in alpha ERD responses in the context of reward feedback processing. Therefore, in the current study we investigated the extent to which alpha ERD is reduced in patients with SZ engaged in a reward task, and importantly how these deficits may relate to negative symptoms.

The majority of prior neuroimaging reward studies in schizophrenia model reward processing in contexts that demand decision-making and/or motor responding to obtain reward. Though this prior literature has made valuable contributions to characterizing schizophrenia-related reward deficits, the predominant use of performance-dependent reward tasks confounds isolation of basic reward responses from processes related to higher-order functions supporting reward attainment. Previous studies have shown that neuro-oscillatory features are modulated during slot machine play in healthy individuals(29, 30). In the current study, we investigate reward-attention interactions by using a slot machine task, in which reward outcome did not depend on participant input, as an ecologically valid approach to model monetary reward processing in the absence of performance demands. Specifically, we analyzed reward outcome processing by assessing post-reward outcome alpha ERD responses to equiprobabe slot wins and final-reel losses.

Prior work from our group using this task did not focus on alpha band, but instead established expected condition effects in heathy controls, including outcome modulations of oscillatory power in response to wins (delta) and losses (theta) (31); and also time-domain event-related potential comparisons between SZ and healthy control (HC) groups that revealed intact anticipatory (stimulus preceding negativity) and early outcome (reward positivity), with late in outcome processing (late positive potential)(32). In the current study, we hypothesized that alpha ERD would be deficient during reward processing in SZ (i.e., less alpha suppression to wins, relative to near miss losses), and that ERD deficits would correlate with negative symptoms. Further, as baseline alpha oscillations are thought to reflect a state of internal rest and neural binding across the brain (referred to as the inhibition hypothesis), with ERD reflecting “desynchronization” of this rest state to ready the brain for task engagement (12), we further hypothesized that trait rumination, reflective of excessive internal focus(33), would also relate to reduced alpha ERD.

## Methods

### Participant Recruitment and Demographics

Study participants were 54 SZ (75.9% men; age range = 19.07 – 64.70 years) and 54 HC (79.6% men; age range = 19.25 – 64.41 years) recruited via community advertisements and from local medical centers. The Structured Clinical Interview for DSM-IV (SCID-IV-TR) (34) was used to confirm a schizophrenia or schizoaffective diagnosis for SZ subjects and to exclude HC if they met criteria for a past or current Axis I disorder. HC were also excluded for having a first-degree relative with a schizophrenia-spectrum disorder. Urine toxicology tested for common drugs of abuse (e.g., opiates, cocaine, amphetamines) and potential subjects with a positive test were excluded. English fluency was required for participation. Additional exclusion criteria for SZ and HC were history of head injury, neurological illness, or other major medical illness that impacts the central nervous system. Written informed consent was obtained from all study participants under protocols approved by the University of California, San Francisco Institutional Review Board.

### Clinical symptom and trait rumination assessment

Negative symptoms were evaluated in the SZ group using the Positive and Negative Syndrome Scale (PANSS (cite). In addition to a negative symptom total score, we also used the PANSS to compute a Depression composite score based on a previously validated five-factor model shown to correlate with depression in SZ (35), similar to our previous work (32); the composite is derived by summing the anxiety (G2), guilt (G3), and depression (G6) items from the general psychopathology subscale. Trait rumination was assessed in both the HC and SZ groups with the Rumination Response Scale (RRS) (36). Symptom ratings interviews were held within one week of EEG recording.

### EEG Data Acquisition and Processing

EEG data were recorded from 64 channels using a BioSemi ActiveTwo system (www.biosemi.com). Electrodes placed at the outer canthi of both eyes, and above and below the right eye, were used to record vertical and horizontal electro-oculogram (EOG) data. EEG data were continuously digitized at 1024Hz and referenced offline to averaged mastoid electrodes before applying a 0.1Hz high-pass filter using ERPlab (37). Data were next subjected to Fully Automated Statistical Thresholding for EEG artifact Rejection (FASTER) using a freely distributed toolbox (38). The method employs multiple descriptive measures to search for statistical outliers (> ±3 SD from mean): (1) outlier channels were identified and replaced with interpolated values in continuous data, (2) outlier epochs were removed from participants’ single trial set, (3) spatial independent components analysis was applied to remaining trials, outlier components were identified using the ADJUST procedure(39), data were back-projected without these components, and (4) outlier channels were removed and interpolated within an epoch. The original FASTER processing approach was modified between steps 2 and 3 to include canonical correlation analysis (CCA). CCA was used as a blind source separation technique to remove broadband or electromyographic noise from single trial EEG data, generating de-noised EEG epochs. This approach is identical to the CCA method described in our prior reports (40, 41).

### Time-frequency analysis (TFA)

Time frequency analysis of single trial EEG data was done with Morlet wavelet decomposition using FieldTrip software (42) in Matlab. Specifically, we used a Morlet wavelet with a Gaussian shape defined by a ratio (σ_f_ = f/C) and 6σ_t_ duration (f is the center frequency and σ_t_ = 1/(2πσ_f_). C was set to seven. In this approach, as the frequency (f) increases, the spectral bandwidth (6σf) increases. As such, center frequencies were set to minimize spectral overlap, resulting in 10 frequency bins: 3, 5, 7, 10, 14, 20, 28, 40, 56, and 79 Hz. The alpha frequency was of particular interest and had a spectral bandwidth of 7.86 to 12.14Hz. Total power was calculated by applying a 10% trimmed mean average to the squared single trial magnitude values in each frequency bin on a millisecond basis. The average power values were 10log_10_ transformed and then baseline corrected by subtracting the mean of the pre-R2 stimulus baseline (−250 to −150 ms) from each time point separately for every frequency. The resulting values describe change in total power relative to baseline in decibels (dB).

### Data Analysis Plan

#### Principal Components Analysis (PCA) for Alpha Power extraction

Similar to our prior work (43, 44), we used a time frequency principal components analysis (TF-PCA)(45) to extract peak event-related spectral power in the alpha range, as it provides an objective data-driven approach to quantifying time-frequency activity in two-dimensional space. Two-dimensional TF data can be subjected to temporal factor analysis approaches by transposing and concatenating these 2D data and treating them like one-dimensional ERP data in the factor analysis (46). The data-driven TF-PCA approach is beneficial because other approaches to quantifying TF activity in 2D space often involves averaging within some subjective temporal and spectral window. To minimize the number of columns (variables) in the TF-PCA, samples approximately every 8ms from 1550ms before R3 onset to 950ms after it were taken for each of the 10 frequencies in the TF matrices. All subjects (n=108), electrodes (n=64), and task conditions (n=3: R3-win, R3-near miss loss, R2-total miss loss) were rows (observations) in the TF-PCA. Total miss loss trials were down-sampled by 50% to match the NM and win conditions. The TF matrices for baseline-corrected total power were submitted to independent, unrestricted covariance-based PCA with promax rotation applied. The PCA was implemented in Matlab (45, 47). All components were retained and subjected to a varimax and then promax rotation, yielding oblique factors corresponding to major TF components. To produce interpretable TF factor loadings, 1-D factor loadings were rearranged back into the original order of the 2-D TF measures.

The primary factor fell within the alpha range (10Hz) peaking at 446 ms (FWHM: −140 to 813ms) post-outcome and accounting for 9.9% of the total variance; (Figure 2a). Standardized, unitless factor scores derived from this primary factor were extracted, per electrode and subject, and subjected to further statistical analysis, as describe below.

**Figure 1.**
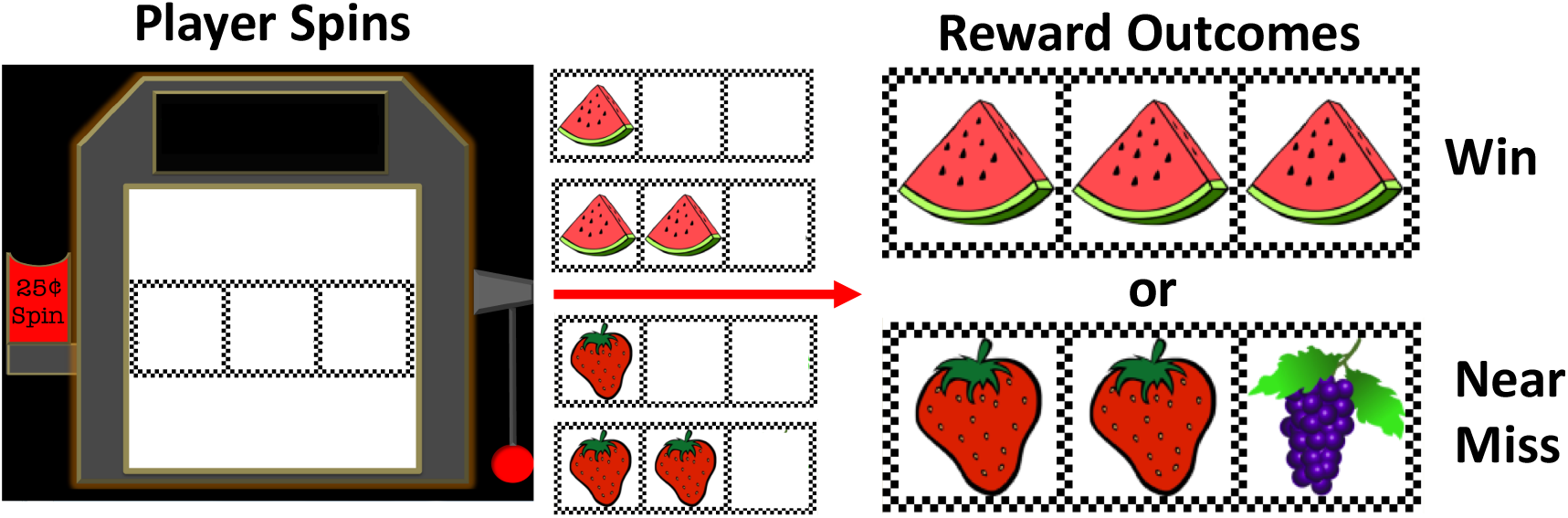
Slot machine task illustration depicting win and near miss loss outcomes revealed at the final (3^rd^) reel.

**Figure 2.**
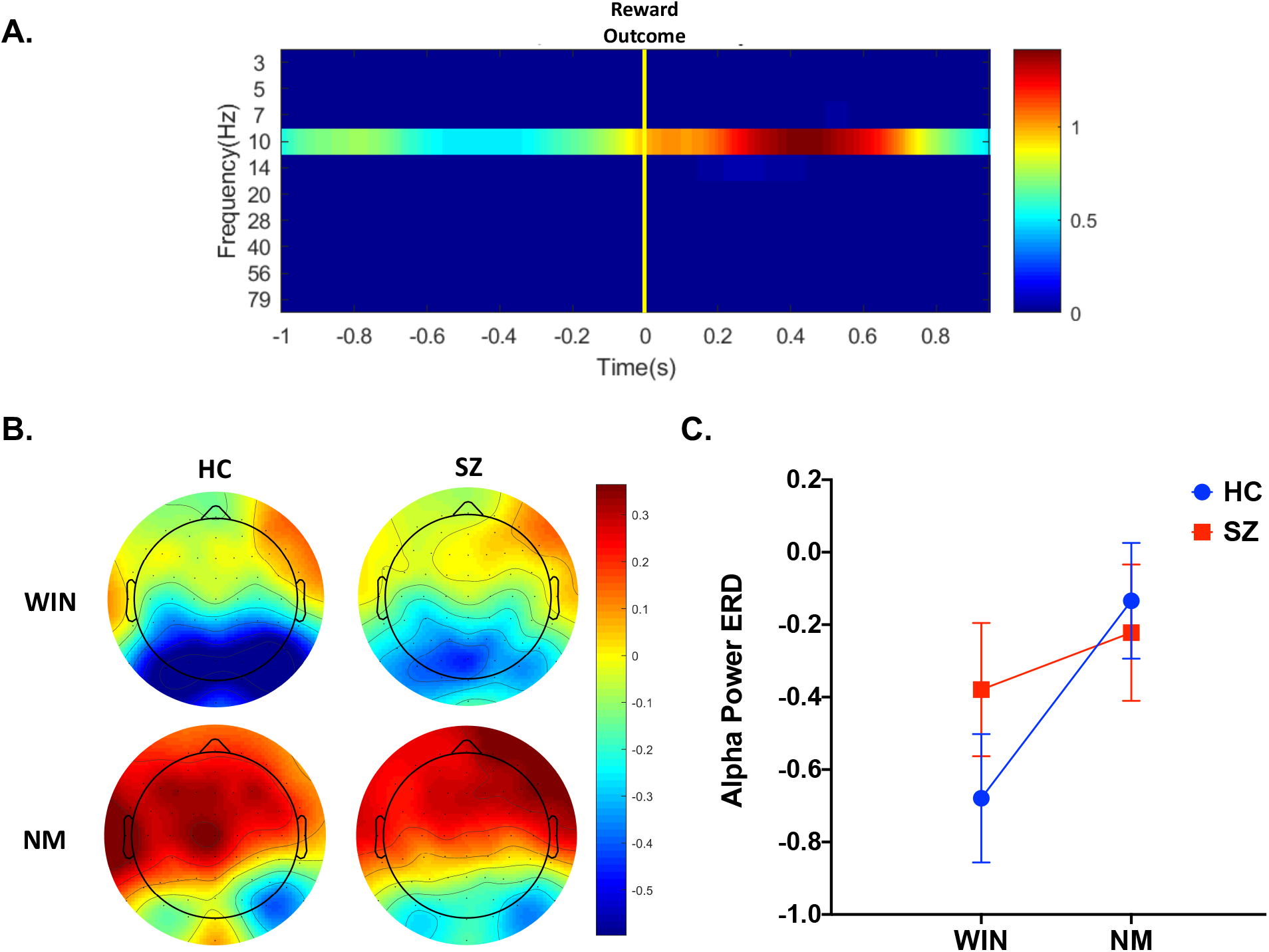
(A, top): The loading plot shows the primary factor derived from the Promax-rotated time-frequency factor analysis of total power, showing an alpha band event-related power component centered at 10 Hz that peaks ~450 ms after reward outcome. Time 0 s (depicted with yellow vertical line) corresponds to the Reel-3 reward outcome reveal. (B, bottom left): Within group average scalp topography maps for alpha ERD factor scores are plotted for healthy control (HC) and schizophrenia (SZ) groups for winning (WIN) and near miss loss (NM) outcomes. (C, bottom right): Mean +/− SE line graphs of total power factor scores averaged over eight posterior-occipital sites displaying WIN vs. NM loss difference effects for each group.

#### Alpha ERD Group effects

Posterior-occipital alpha total power factor scores were modeled with repeated-measures Analysis of Variance, with Group as a between-subject factor (HC, SZ), and within-subject factors for Reward Outcome (win, near miss loss) and Electrode Site (PO7, POZ, PO8, PO3, POZ, PO4, O1, OZ, O2). The omnibus effect of interest was the 2-way Group by Reward Outcome, which, if significant (p<.05), was followed-up with paired t-test within each group (follow-up testing was Bonferroni-corrected across 2 levels of group; p<.025).

#### Alpha ERD relationships with clinical symptoms and trait rumination

In order to examine correlations with clinical variables, ERD was averaged over the 8 posterior-occipital electrode sites to create a single posterior alpha ERD value for each individual, for each condition (win, near miss loss). Correlations of i) symptoms within the SZ group, and ii) trait rumination among all participants was examined using multiple regression to test for differences in the regression line slopes between the HC and SZ groups, and if the slopes did not significantly differ, to test the pooled estimate of the common slope of the regression line for significance. These tests were Bonferroni-corrected across 2 levels of condition; p<.025. Data from 3 SZ individuals were missing on the RRS rumination measure, and so these analyses were undertaken in n=54 HC; n=51 SZ).

Lastly, within the SZ group, relationships with negative symptoms, antipsychotic medication dosage and illness duration were examined and similarly Bonferroni-corrected across 2 levels of condition; p<.025. The continuous duration of illness variable was log-transformed to mitigate right skewness. Distributions of all other variables were examined and were suitably distributed for parametric statistics without transformation.

## Results

### Posterior-Occipital Alpha Power ERD Group effects

In the rmANOVA model examining group alpha ERD to reward outcomes, we observed a significant Group by Reward Outcome interaction effect (F(1,106) =5.77, p=.018). Planned, Bonferroni-corrected follow-up testing at each level of Group was then conducted to parse the interaction effect. Within HC, winning outcomes elicited a significantly stronger alpha power ERD to wins than near miss losses (t(53) = −4.71, p<.001). In contrast, SZ showed no alpha ERD differences in response to win vs. near miss loss outcomes (t(53) = −1.40, p=.167). See Figure 2b and 2c.

### Trait Rumination relationships with Alpha ERD, in HC and SZ groups

First, the SZ group endorsed higher trait rumination, relative to the HC group (F(1,103) = 28.92, p<.001). Multiple regression was then used to test for group differences in regression slopes relating self-reported rumination to average posterior alpha ERD. Slope difference models showed no interaction effect that would indicate a significant group differences in the slopes relating rumination and alpha ERD (wins F(3,102) = 1.30, p = .257; near miss losses F (3,102) = 2.00, p=.260). Therefore, the group interaction terms were dropped from these regression models in order to test for a common slope across both HC and SZ groups. The reduced models revealed a common slope for both win and near miss conditions, indicating that higher trait rumination was associated with significantly less posterior alpha ERD, for both wins and near miss loss conditions (wins F (3,101) = 14.14, p=.005; near miss losses F (3,101) = 9.68, p=.002. When the SZ participant with the highest trait rumination (>3 S.D. from group mean) was removed from these analyses, the associations between rumination and alpha ERD remained statistically significant for both win and near miss loss common slopes. See Figure 3.

**Figure 3.**
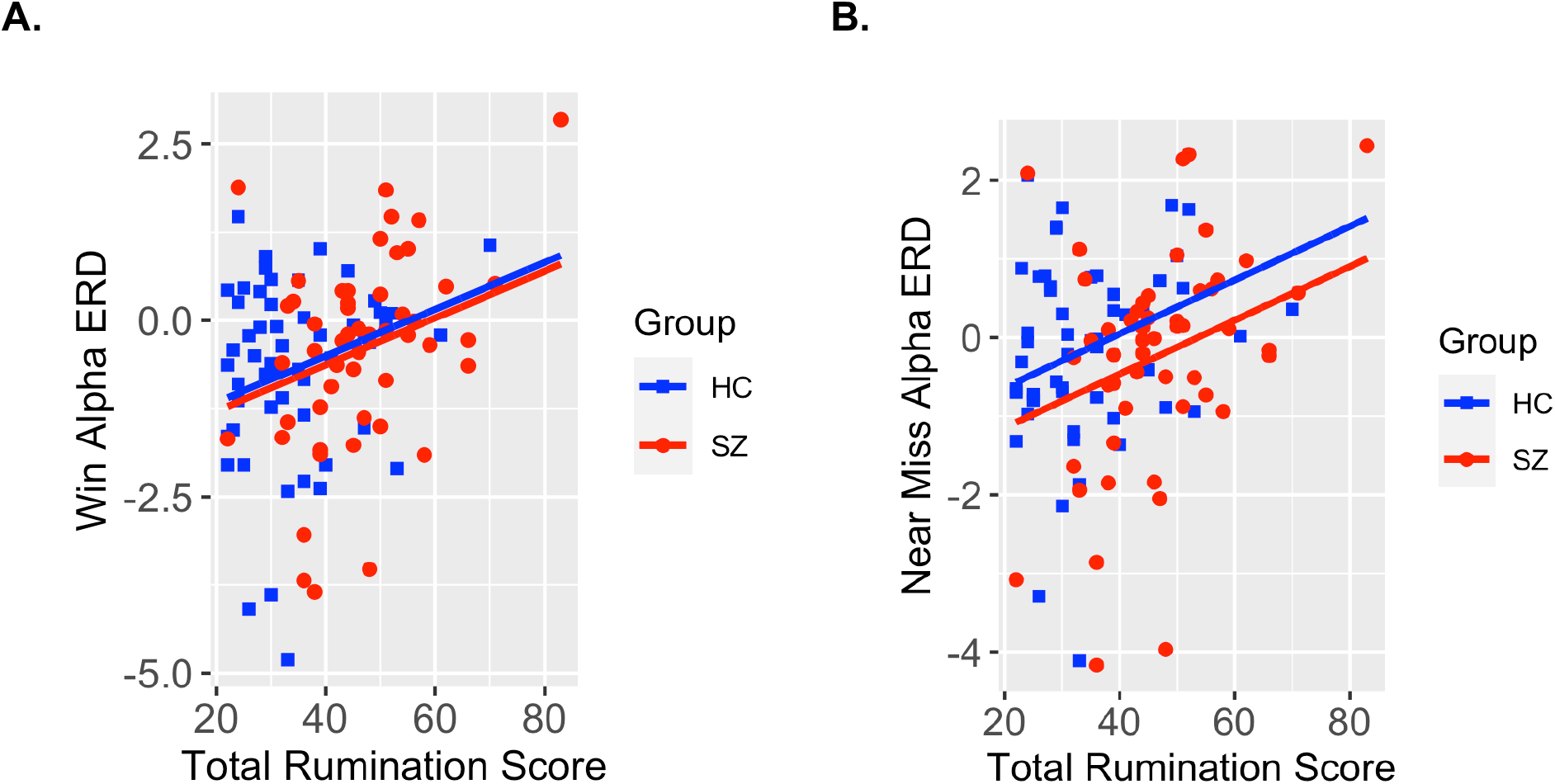
Scatterplots showing slope relationships between trait rumination levels and alpha power factor scores for the healthy control (HC, shown in blue) and schizophrenia (SZ, shown in red) groups for win (left) and near miss loss (right) outcome conditions. P-values correspond to a common slopes regression model as non-significant group interaction term from each model.

### Clinical feature relationships with Alpha Power ERD, within the SZ Group

Within the SZ group, alpha ERD to either outcome type did not correlate with PANSS negative symptoms (p>.163). Nor did alpha ERD correlate with antipsychotic medication dose (p>.456), or with illness duration (p> .413).

## Discussion

The goal of this study was to examine reward and attention system interactions in schizophrenia by assessing alpha ERD in the context of a rewarded slot machine task, and to further relate hypothesized reductions in event-related alpha signaling to both negative symptoms and trait rumination. We observed task-related alpha band power modulations across distributed bilateral posterior scalp electrode sites, peaking ~450 ms after reward outcome information became available. The principal finding of this study was a prominent difference in HC between alpha ERD to wins vs. losses which was not present in SZ, suggesting that modulation of attention related to alpha is deficient in schizophrenia during reward feedback processing.

First, we observed significant stimulus-locked alpha desynchronization in HCs to wins compared to near miss losses, consistent with alpha ERD’s canonical role in “task engagement” and attention. Of note, while prior HC studies have largely focused on the role of other oscillatory bands (such as theta) in examining reward processing, here we show significant differences in the degree or alpha ERD between the wins and near-miss trials in our HC group; this extends understanding of alpha ERD’s role in attentional engagement to include interaction with reward system processes. Two prior studies from the same research group did not show changes in alpha (or theta) power elicited by near miss (vs. win) events (29, 30), though both were based on a slot task in which near misses occurred with twice the frequency as wins, making it difficult to compare results with our study in which trial numbers were balanced across near miss and win outcomes to promote disentangling brain responses related to outcome processing from probability effects. While prior studies examining alpha oscillations in reward have reported enhancement of alpha ERD during reward anticipation (i.e. in the period just prior to reward receipt) (reviewed by(10)), our findings point to reward-outcome specific effects (i.e. elicited by reward receipt) on alpha oscillatory power, and that both reward anticipation and reward receipt may exert modulatory effects on higher order cognitive processes via alpha ERD.

Next, we found attenuated reward alpha ERD, (i.e., less alpha suppression to wins, relative to near miss losses in the SZ group compared to the HC pattern) suggesting deficient reward system modulation of attention and task engagement in patients with schizophrenia. Between-group alpha ERD effects may mean that HC deploy more visual attentional resources to wins than losses, whereas alpha signaling in SZ more weakly distinguishes reward from non-reward outcomes. The majority of neuroimaging studies of reward processing in schizophrenia have relied on tasks that require decision making, response planning and response execution to obtain rewards. Therefore, less is known about whether reward processing deficits endure in schizophrenia in the absence of these higher-order response. It is therefore worth emphasizing that given our relatively passive slot machine task design, in which reward outcome did not depend on participant performance or decision-making, we observe alpha ERD alterations in schizophrenia even when reward attainment places minimal demands on higher-order cognitive processes. This finding is of particular interest as our group has previously shown intact reward anticipation and early outcome reward processing in this same experimental cohort(32); event-related potential analyses in our group’s earlier work demonstrated no differences between SZ and HC in either the stimulus preceding negativity (SPN), a pre-stimulus onset ERP reflecting reward anticipation, or the reward positivity (RewP), an early (~250ms) evolving ERP reflecting early reward feedback processing. This intact RewP is in line with prior studies examining feedback processing in SZ and supports current conceptualizations that patients with schizophrenia have intact hedonic responses. In contrast, our findings demonstrate altered neural synchrony, specific to alpha activities during outcome evaluation, suggesting that modulation of attention and information processing related to alpha is deficient in schizophrenia during reward processing. Our current findings therefore suggest that reward deficits in schizophrenia may be better reflected in later stage reward outcome processing, including attenuated reward system modulation of attentional processing, which in turn may contribute to the motivational and avolitional deficits widely described in schizophrenia.

We did find a significant relationship between trait rumination and alpha ERD in our combined HC and SZ sample. Rumination, defined as repetitive thoughts focused on negative or worrying content(48), occurs in both neurotypicals and across a number of psychiatric diagnoses including in depression and many anxiety disorders(33, 48, 49). Further, rumination severity is associated with worsened psychiatric morbidity, such as longer and more severe depressive episodes(50). Prior studies have shown that excessive rumination impairs engagement of cognitive processes comprising executive functioning including attention, memory retrieval and cognitive flexibility(51–53), and is further reflected in task-relevant reductions in oscillatory power, including in the alpha and beta bands (49). Rumination has been described as a negative internally focused state which if excessive can impair engagement of externally focused cognitive processes such as attention(54). Indeed, this conceptualization of aberrant “internal” focus is congruent with work describing excessive default mode network (DMN) functional connectivity as a core network mechanism in depression (55) and psychosis (56). Our findings associating greater rumination with relatively attenuated alpha ERD are also consistent with these explanatory frameworks describing impaired relationships between “internal” and “external” focus. As alpha oscillations are thought to reflect an “at rest” or “idle” brain state (during which rumination would likely be most active), it is possible that stronger ruminative tendencies could reflect a more static “internal” resting state leading to difficulty in transitioning to an “on-line” task engaged state, as reflected in the alpha ERD deficits we report here.

In the current study we did not demonstrate significant relationships between attenuated alpha ERD and negative symptoms in the SZ cohort as we had hypothesized. It is possible that alpha ERD truly has no relationship to negative symptomology, and that other neurophysiological mechanisms related to the reward system explain negative symptom variance. Alternatively, there are several other reasons that our study could have failed to observe a relationship between alpha ERD and negative symptoms, including general reasons such as measurement assessment windows and psychometrics, e.g., (57), as well as study specific explanations, such as we did not oversample for negative symptoms or deficit syndrome. Future studies focusing on oscillatory phase synchronization (e.g., inter-trial coherence measures) in addition to magnitude of neuro-oscillatory responses will be important in extending this work. Further, we chose to focus on oscillatory measures within the alpha range, given our interest in focusing on attention and information processing functions ascribed to alpha ERD. Yet it is important to acknowledge that other frequency bands have been implicated in schizophrenia pathology, including gamma band oscillations and their role in local glutamatergic circuits that support cognition. It is therefore worth noting that through cross-frequency phase and amplitude coupling mechanisms in which lower frequencies gate higher ones (11), alpha activities may also exert influence on neural processes oscillating at higher frequency ranges, like gamma. To fully characterize electrophysiological dynamics relevant to reward - attention system interactions, more comprehensive assessments including other frequency bands are needed, and well as examination of their coordination through cross-frequency coupling (58). The current study linking reward-outcome to alpha ERD provides an important step in elaborating the complex interplay between reward and attentional systems, and how the reciprocal interactions between these two systems may break down in schizophrenia.

**Table 1.**
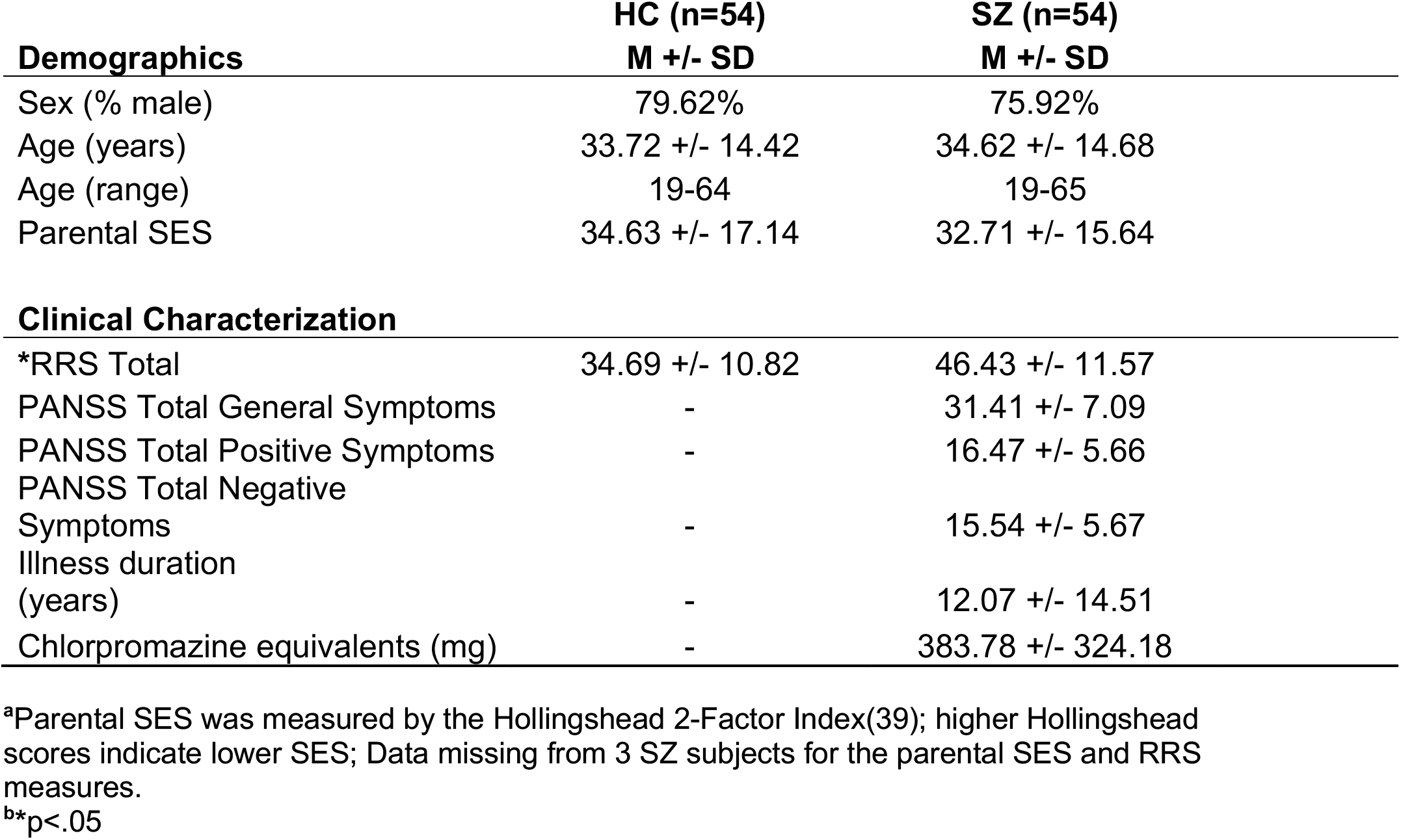
Sample demographics and clinical variables.

|  | **HC (n=54)** | **SZ (n=54)** |
| --- | --- | --- |
| **Demographics** | **M +/- SD** | **M +/- SD** |
| Sex (% male) | 79.62% | 75.92% |
| Age (years) | 33.72 +/- 14.42 | 34.62 +/- 14.68 |
| Age (range) | 19-64 | 19-65 |
| Parental SES | 34.63 +/- 17.14 | 32.71 +/- 15.64 |
| **Clinical Characterization** |  |  |
| *RRS Total | 34.69 +/- 10.82 | 46.43 +/- 11.57 |
| PANSS Total General Symptoms | - | 31.41 +/- 7.09 |
| PANSS Total Positive Symptoms | - | 16.47 +/- 5.66 |
| PANSS Total Negative Symptoms | - | 15.54 +/- 5.67 |
| Illness duration (years) | - | 12.07 +/- 14.51 |
| Chlorpromazine equivalents (mg) | - | 383.78 +/- 324.18 |

<sup>a</sup>Parental SES was measured by the Hollingshead 2-Factor Index(39); higher Hollingshead scores indicate lower SES; Data missing from 3 SZ subjects for the parental SES and RRS measures.
<sup>b</sup>\*p<.05

## Acknowledgments

Research supported by VA CX001028 and CX001980 to Dr. Fryer. Dr. Ford is supported by a VA Senior Research Career Scientist award. Abagail Blaine assisted with manuscript preparation and data curation.

